# Resting state fMRI reveals differential effects of glucose administration on central appetite signalling in young and old adults

**DOI:** 10.1101/529552

**Authors:** Riccarda Peters, David J White, Andrew Scholey

## Abstract

**Background:** Healthy aging has been associated with reduced appetite and energy intake, which can lead to loss of bodyweight and undernutrition and related health problems. The causes for the decline in caloric intake are multifactorial involving physiological and non-physiological processes.

**Aims:** Here we examined age-related, physiological changes in brain responses associated with macronutrient intake.

**Methods:** Using a randomized, double blind, balanced cross-over design, younger (n=16, aged 21-30) and older adults (n=16, aged 55-78) received a drink containing glucose and a taste-matched placebo after an overnight fast. Blood glucose and hunger were assessed at baseline and 20 minutes post-ingestion, after which participants underwent resting state functional magnetic resonance imaging (rsfMRI).

**Results:** Frequency dependent changes in slow-5 (0.01-0.027Hz) and slow-4 (0.027-0.073Hz) amplitude of low frequency fluctuations (ALFF) and fractional ALFF (fALFF) of the blood oxygen level-dependent (BOLD) signal were contrasted between sessions and age groups. We observed a significant treatment x age-group interaction in slow-5 ALFF and fALFF in the left insula. Younger participants showed a decrease in BOLD amplitude, whereas older participants showed an increase. We further observed a treatment x age-group interaction in slow-4 ALFF in the occipital and lingual gyrus and precuneus with older participants showing an increase in magnitude of slow-4 ALFF and younger participants showing a decrease in the same measure.

**Conclusion:** These age-related, frequency-dependent changes in the magnitude of the BOLD signal in a key region related to energy homeostasis following feeding may contribute to behavioral changes in energy intake during senescence.

## Introduction

A recent meta-analysis reported that healthy older adults (aged 70-74) have a 16-20% lower energy intake and 25-39% lower hunger than younger adults (aged 26-27) (Giezenaar et al., 2016). Reduced nutrient and energy intake may increase the occurrence of undernutrition (Mowe et al., 1994). Low body weight and loss of lean tissue are strong predictors of poor health outcomes including those that contribute to premature death (Landi et al., 2016; Thibault et al., 2011). The decline in caloric intake with increasing age is referred to as anorexia of aging (Morley, 1997). Evidence suggests that the development of anorexia of aging is multifactorial, including both physiological impairments of food intake regulation and non-physiological (e.g. social and psychological) processes (Hays and Roberts, 2006; de Boer et al., 2013).

Unravelling the physiological age-related changes in brain response to energy intake may aid in understanding pathogenesis of anorexia of aging. The central regulation of appetite in healthy individuals is a complex system resulting from an interplay between neurochemical compounds and signaling from homeostatic neuronal circuits (Schwartz et al., 2000). The use of functional magnetic resonance imaging (fMRI) has been helpful in identifying brain structures involved in the regulation of energy intake. A key region in homeostatic metabolic regulation is the hypothalamus. Several studies have demonstrated that hypothalamic blood oxygen level-dependent (BOLD) signal decreases after glucose administration (Smeets et al., 2005; Smeets et al., 2007; Little et al., 2014). Other studies have shown that resting state functional connectivity of hypothalamus and insula are altered in response to changes in homeostatic energy balance (Wright et al., 2016). While these studies focused on specific regions of interest, more recent studies have used data-driven approaches to investigate changes in caloric intake associated with global brain homeostasis. A recent study compared whole brain BOLD intensities before and after glucose ingestion and reported decreased BOLD activity in brain regions associated with reward and feeding including the insula, thalamus, anterior cingulate, orbitofrontal cortex, amygdala and hippocampus (van Opstal et al., 2018).

Human brain function constitutes complex and dynamic systems that generate a myriad of neuro-oscillatory rhythms. These underpin correlated activity between brain regions including resting state networks. The indices of the amplitude of low frequency fluctuations (ALFF) (0.01-0.1 Hz) (Zang et al., 2007) and fractional amplitude of low frequency fluctuations (fALFF) (Zou et al., 2008) are related measures that quantify the amplitude of these low-frequency oscillations in the BOLD signal. A recent study identified the fractional amplitude of low frequency fluctuations (fALFF) as a sensitive measure to detect homeostatic changes in the resting brain (Al-Zubaidi et al., 2018).

ALFF measures the intensity of regional spontaneous brain activity and is defined as the total power within the frequency range (0.01-0.1 Hz), while fALFF measures the relative contribution of low-frequency fluctuations and is defined as the power within the low-frequency range (0.01-0.1 Hz) divided by the total power in the entire frequency range (Zuo et al., 2010). Both measures have high temporal stability (Küblböck et al., 2014) and test-retest reliability (Zuo and Xing, 2014). While fALFF has been reported to have greater specificity to grey matter, ALFF has been reported to have higher test-retest reliability (Zuo et al., 2010). Therefore, ALFF and fALFF are often reported together to maximise reliability across subjects.

In more comprehensive investigations of amplitude information of low frequency oscillations, the power spectra of rsfMRI has been decomposed into different frequency bands. Subsets of frequency bands have been reported at slow-5 ( 0.01-0.027 Hz), slow-4 (0.027-0.073 Hz), slow-3 (0.073-0.198 Hz) and slow-2 (0.198-0.25Hz) (Zuo et al., 2010). These distinct infra-slow frequency ranges have been proposed to be generated by distinct oscillators, each associated with distinct characteristics and physiological functions. Grey matter related oscillations have been shown to primarily occur in the slow-4 and slow-5 range (0.01-0.073 Hz), while slow-3 and slow-2 signals occurred in white matter (Zuo et al., 2010).

To our knowledge, no study has investigated frequency-specific changes in ALFF and fALFF as a function of energy intake. The aims of this study were twofold, (i) to examine frequency dependent changes in ALFF and fALFF in the slow-5 and slow-4 frequency window in response to glucose ingestion, and (ii) to determine whether older participants showed abnormalities in the response to glucose fluctuations in specific frequency bands. It was hypothesised that there would be differences in ALFF and/or fALFF in different frequency bands in homeostatic brain regions between young and older participants. We further hypothesised changes in ALFF/fALFF magnitude would be related to self-perceived hunger ratings.

## Method

### Subject Characteristics

Sixteen younger participants (8 female, aged 21-30) and sixteen older participants (8 female, aged 55-78) were recruited through local advertising and from a database. All participants had normal or corrected-to normal vision and hearing, no major physical illness and had no history of neurological or psychiatric illness or head trauma. Further exclusion criteria were a diagnosis of diabetes mellitus, a history of hypersensitivity to glucose, heart disease or high blood pressure, smoking, substance abuse, intolerance to artificial sweeteners, pregnancy, claustrophobia, metal implants or any other contraindications to MRI.

Participants were excluded if they reported health conditions that would affect food metabolism including the following: food allergies, kidney disease, liver disease and/or gastrointestinal diseases (e.g. irritable bowel syndrome, coeliac disease, peptic ulcers). Subjects were excluded if they were taking any medication, herbal extracts, vitamin supplements or illicit drugs which might reasonably be expected to interfere with blood glucose levels within four weeks prior to and during the study.

This study was approved by the Swinburne University Ethics committee and all study procedures were performed in accordance with the principles of the 1964 Declaration of Helsinki. All subjects provided written informed consent.

### Study design

Participants attended one screening/familiarisation session and two experimental sessions for this randomised, double-blind, crossover study. All testing took place at Swinburne University of Technology, Melbourne. The two testing sessions were scheduled to be at least two days but not more than two weeks apart and started between 8.30am and 10am for all participants. Treatment assignment followed a Latin square design so that order of treatments was counterbalanced within age groups and gender. To ensure that the active and placebo treatment were indistinguishable in taste (Scholey et al., 2001) the vehicle was 150ml of water mixed with 20 ml sugar free cordial. For the glucose condition 25 g glucose (Glucodin Pure Glucose Powder) were dissolved in the vehicle. For the placebo visit the vehicle was mixed with two tablets (30 mg) sodium saccharine.

Blood glucose levels were measured at baseline, 20 min post-ingestion, 120 min post-ingestion and 150 min post-ingestion. For the purpose of the present analysis we will focus on the baseline and 20 min post-ingestion measures. Glucose was measured via capillary fingerprick using a Freestyle Optium Blood Glucose Sensor and Optium Blood Glucose Test Strips (Abbott Diabetes Care Ltd., Witney, UK).

Treatment preparation and assignment and blood glucose measurements were performed by a disinterested third party not further involved in the study. Resting state data and blood glucose levels of this study have been explored under a separate hypothesis relating glucose modulation of hippocampal connectivity and are reported elsewhere (Peters et al., 2018).

### Hunger Rating

Subjective feelings of hunger were assessed on a computerised Visual Analogue Scale (VAS) before each blood sample. Participants were shown a straight 100 mm line with the endpoints labelled “not at all hungry” and “extremely hungry” respectively. Subjects were instructed verbally to rate on the line how hungry they felt in the present moment and to avoid the extreme ends of the line, as this would indicate the least and most hungry they have ever felt in their life respectively. Thirst, attention and overall mood were also assessed with VAS and are reported in supplementary material (S1.1).

### MRI acquisition

MRI scanning was performed on a 3-Tesla Siemens Magnetom Trio scanner (Siemens, Erlangen, Germany) with a 32-channel head coil. Anatomical high resolution 3D T1-weighted magnetization prepared rapid acquisition gradient echo (MP-RAGE) were acquired (1mm isotropic MP-RAGE, TR= 1900ms, TE = 2.52ms, flip angle= 9°). Resting state scans were acquired with an interleaved multi band sequence (multiband acceleration factor= 6, bandwidth= 1860 Hz/Px, TR= 870ms, TE= 30ms, echo spacing= 0.69ms, flip angle= 55°, field of view= 192mm, voxel resolution= 2×2×2mm, slice orientation= transversal, number of slices= 66, 500 volumes per session, total acquisition time: 7mins 15sec).

### Preprocessing

Preprocessing of structural and functional images was conducted using Statistical Parametric Software (SPM) (SPM12; Wellcome Department of Imaging Science, Functional Imaging Laboratory, University College London) and the functional connectivity toolbox CONN V17f (http://www.nitrc.org/projects/conn) (Whitfield-Gabrieli and Nieto-Castanon, 2012) run under Matlab R2014a.

Functional images were first realigned and unwarped, then ART-based outlier identification was conducted. An image was identified as outlier if composite movement from preceding image exceeded 0.5mm or if global mean intensity was greater than 3 standard deviations from mean image intensity for the run in which it was collected. Images were then simultaneously segmented into grey matter (GM), white matter (WM) and cerebrospinal fluid (CSF) and normalized to MNI space (Montreal Neurological Institute, Canada).

Normalized images were smoothed with a 4mm full width half maximum (FWHM) Gaussian kernel. The aCompCor (Behzadi et al., 2007) approach was used to mitigate confounding effects from the BOLD time series. Specifically, five eigenvectors of the principal component decomposition from WM and CSF (derived via segmentation of anatomical images), as well as 12 motion regressors (six head motion parameters and six first-order temporal derivatives) of realignment and parameters derived from ART outlier identification, were regressed out of the BOLD functional data to reduce effects of head motion and non-neural BOLD fluctuations.

As an additional step to control for motion, Framewise Displacement (FD; maximum total and averages scan-to-scan) was calculated according to Power et al. (Power et al., 2012) between-sessions and between-groups. At both sessions average FD was higher for younger adults compared to older adults (young vs old on placebo: *T*(28)= -2.48, p= 0.021; young vs elderly on glucose: *T*(28)= -2.181, p= 0.041). Further, a significant difference in FD between sessions was observed in the younger group (placebo vs glucose: *T*(13)=-2.602; p= 0.022).

Therefore, average FD was included as a covariate for all analyses at the second level. Further, total grey matter volume was calculated using the Tissue Volumes utility in SPM12, estimated from the segmentation of the T1-weighted 3D-MPRAGE anatomical image using the Unified Segmentation of SPM12 (Ashburner and Friston, 2005), which incorporates the updated tissue probability maps with six classes, for inclusion of grey matter volume as a covariate in subsequent analyses.

### Frequency specific ALFF and fALFF

Using the frequency-decomposition options in the CONN toolbox we created two condition-specific frequency-windows for each visit (placebo and glucose) covering slow-5 (0.01-0.027) and slow-4 (0.027-0.073) ranges. Voxelwise amplitude of low-frequency fluctuations analysis (ALFF) and fractional ALFF (fALFF) was carried out by extracting power spectra via a Fast Fourier Transform and computing the sum of amplitudes in the previously defined separate slow-5 and slow-4 frequency bands for each session. The ALFF measure at each individual voxel represents the root mean square of BOLD signal amplitude after bandpass filtering (Zang et a l., 2007), for the calculation of fractional amplitude fALFF the total amplitude of the low-frequency range was divided by that of the entire frequency range. All analyses were performed at the whole brain level.

### Statistical Analysis

Changes in blood glucose levels and changes in subjective feelings of hunger from baseline to 20 minutes post-ingestion were assessed using repeated measures analysis of variance (rmANOVA) with two within group factors for treatment visit (“Drink”: glucose, placebo) and timepoint sample taken (“Timepoint”: baseline, 20mins post-ingestion) and between groups factor age group (young, old). SPSS software version 25 (Chicago, USA) was used for statistical analysis of blood glucose levels and subjective feelings of hunger and correlations.

To assess age-related changes in slow-5 and slow-4 frequency ranges in response to glucose ingestion, voxel-wise intervention and group effects were assessed with mixed within subject (glucose, placebo) and between subject (younger, older) second level ANOVA models using the partitioned variance approach as implemented in CONN, for each frequency band separately. All significance tests used a cluster-extent FWE-corrected *p*-value < 0.05, obtained at a cluster-defining threshold of *p* < 0.005 (Friston et al., 1994). Anatomical labelling of significant clusters was conducted according to the AAL atlas (Tzourio-Mazoyer et al., 2002). Amplitude values of significant clusters were extracted and plotted *post hoc* to determine directionality of the effect. To assess whether the reported results were driven by differences in grey matter (GM) volume, all analyses were repeated using individual total GM volume as a covariate using a one-sided contrast.

## Results

Two subjects of the young group (one female, one male) were omitted from the analysis, due to inconsistent baseline blood glucose levels, indicative of non-compliance with fasting. The analysis of reported hunger was restricted to 14 participants in each group, due to missing data of two participants in the older group. Demographic details for the sample are provided in Table 1 below. Participants included in the study did not differ significantly in BMI (*P*= 0.224) or average fasting blood glucose levels (*P*= 0.43).

**Table 1:** Demographic and morphometric characteristics of the young and older cohorts. Means (with SD) are shown with *P*-values for age group differences from independent groups *T*-tests.

|  | Younger |  | Older |  | P-value |
| --- | --- | --- | --- | --- | --- |
|  | (N= 14, 7 female) |  | (N= 16, 8 female) |  |  |
|  | Mean | SD | Mean | SD |  |
| Age (years) | 25 | 3.5 | 68.3 | 6.4 |  |
| Current BMI (kg/m2) | 23 | 4.5 | 25 | 4.01 | 0.224 |
| Average fasting blood glucose | 5.5 | 0.44 | 5.7 | 0.69 | 0.43 |

### Blood glucose levels across placebo and glucose visits

Blood glucose levels were measured at baseline and 20 minutes post-dose at each study visit. Fasting blood glucose levels did not differ significantly between groups at either placebo or glucose visit (placebo: *T*(28)= -0.88, *P*= 0.39; glucose: *T*(28)= 0.59, *P*= 0.95). There was a main effect of Drink (*F*(1, 27)=55.27; *P*=0.001, η^2^=0.67), a main effect of Timepoint (F(1, 27)= 55.96, *P* = 0.001, η^2^= 0.67) and a Timepoint x Age group interaction (*F*(1, 27)= 6.6, *P*= 0.004, η^2^=0.27. After the ingestion of the glucose drink, glucose levels of young and older participant increased. At 20 min older participants’ blood glucose levels were significantly higher than younger participants’ (*T*(28)= -2.98, *P*= 0.006) and the glucose condition was significantly higher than placebo (*T*(28)= -8.33, *P* < 0.0001) (**Figure 1**).

**Figure 1:**
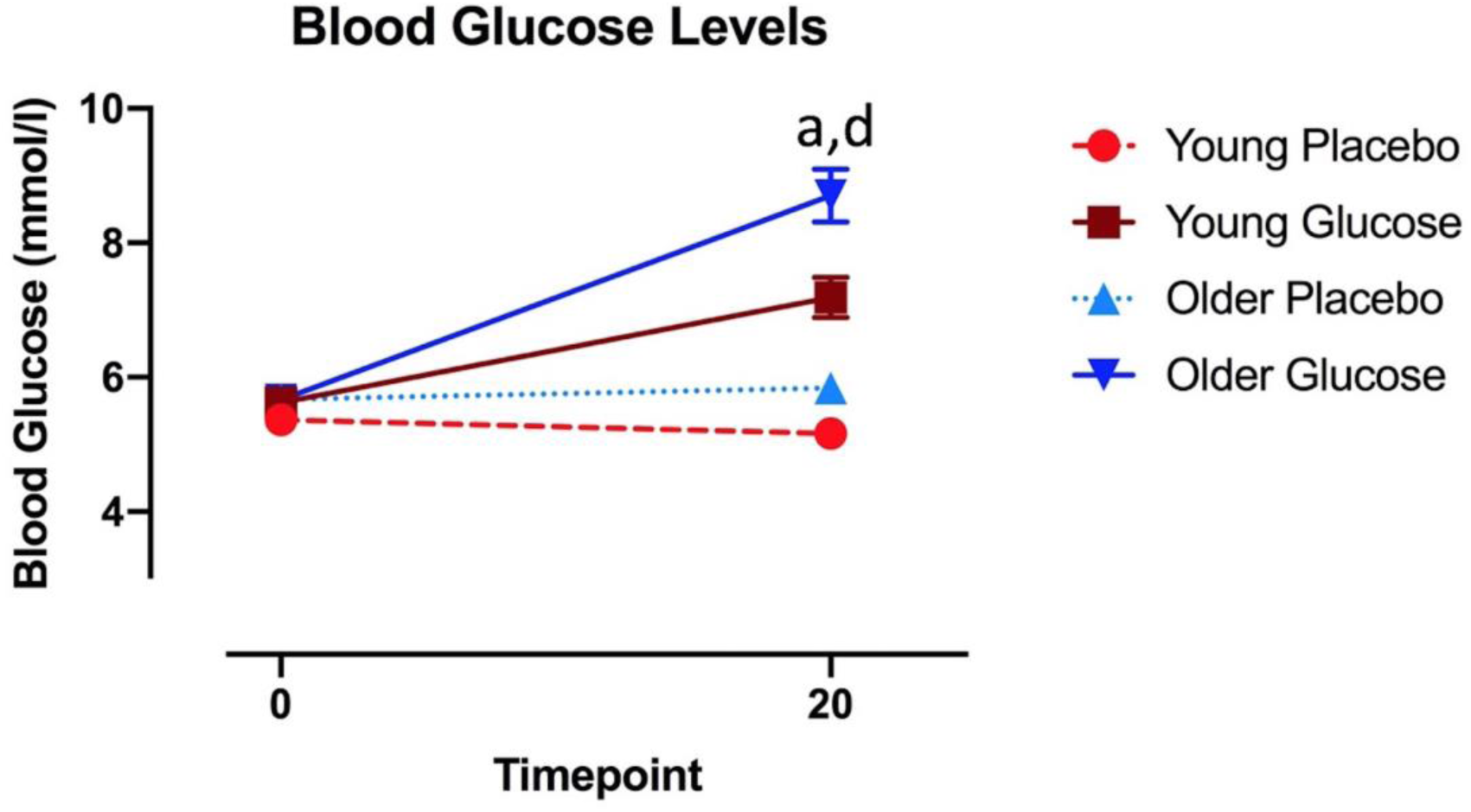
Blood glucose levels. Mean with SEM blood glucose levels at baseline and 20 minutes post-dose at placebo and glucose visit for young (depicted in red) and older group (depicted in blue). Solid lines represent Glucose visit, dotted/dashed lines represent Placebo visit. a= significant age effect and d = significant drink effect at that timepoint.

### VAS Hunger ratings

Hunger ratings were taken immediately prior to blood glucose measurements at baseline and 20 minutes post-ingestion at each visit. Hunger ratings did not significantly differ between young and older adults at any timepoint and there was no Drink x Age group interaction (F(1,26)= 1.64, *P*=0.21, η^2^=0.59). There was however a significant main effect for Timepoint (F(1, 26)= 8.73; *P*= 0.007, η^2^= 0.251). Post-hoc pairwise comparisons revealed that averaged across all participants and across the two treatment types, hunger ratings overall decreased from baseline to 20mins post-ingestion rating (*P*=0.007) (**Figure 2**).

**Figure 2:**
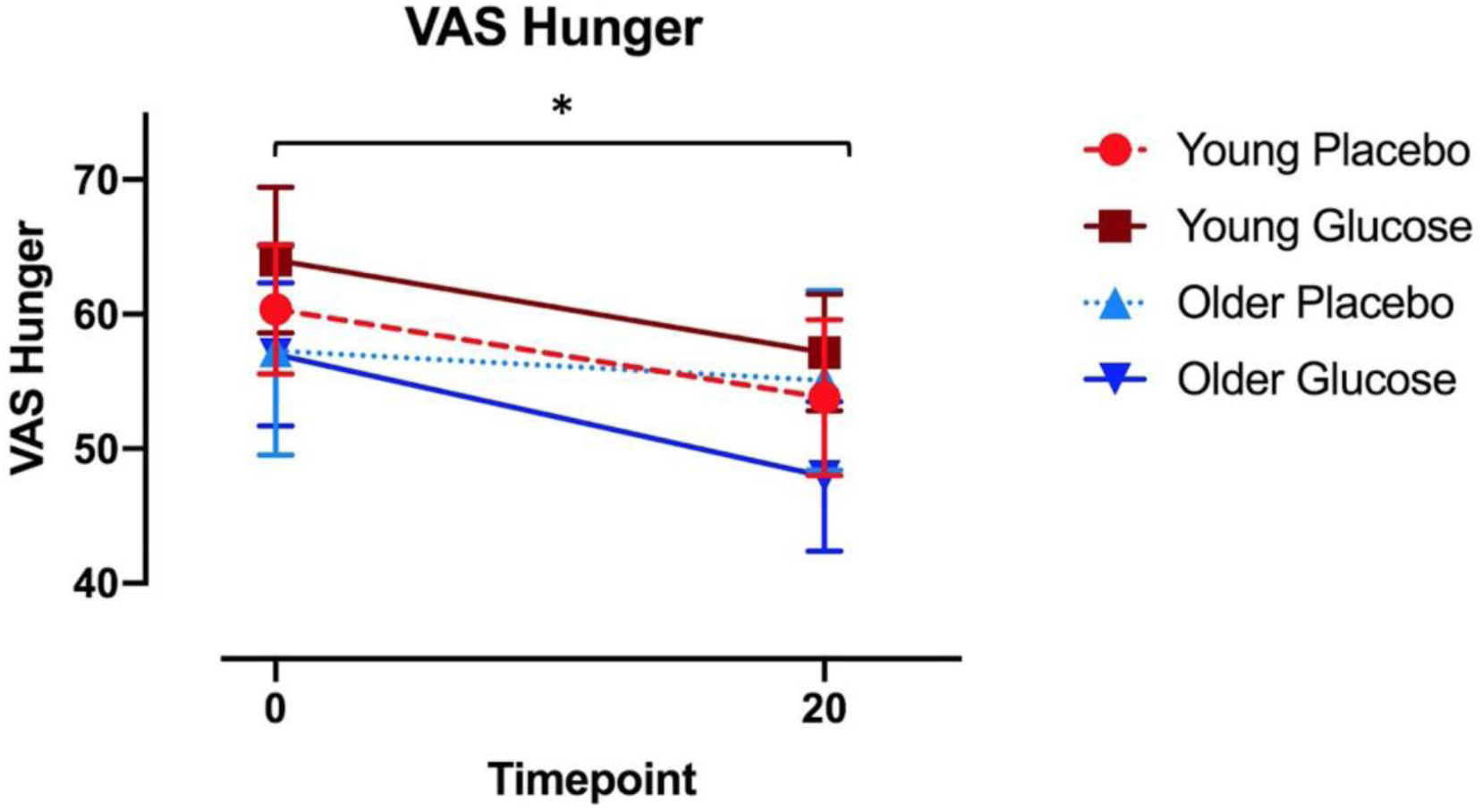
VAS hunger ratings. Mean and SEM VAS Hunger ratings levels at baseline and 20 minutes post-dose dose at placebo and glucose visit for young (depicted in red) and older group (depicted in blue). Solid lines represent Glucose visit, dotted/dashed lines represent Placebo visit. *= main effect of Timepoint.

### Age-related Effect of glucose on BOLD amplitude

#### Slow-5 (0.01-0.027Hz) ALFF: Treatment x age group interaction

We observed Drink x Age group interactions in the left insular cortex (see **Table 2a**; **Figure 3a**). As plotted in **Figure 3b,** ALFF in the left insular cortex decreased after glucose ingestion compared to placebo for the younger adults, while ALFF increased in older adults after glucose administration.

**Table 2:**
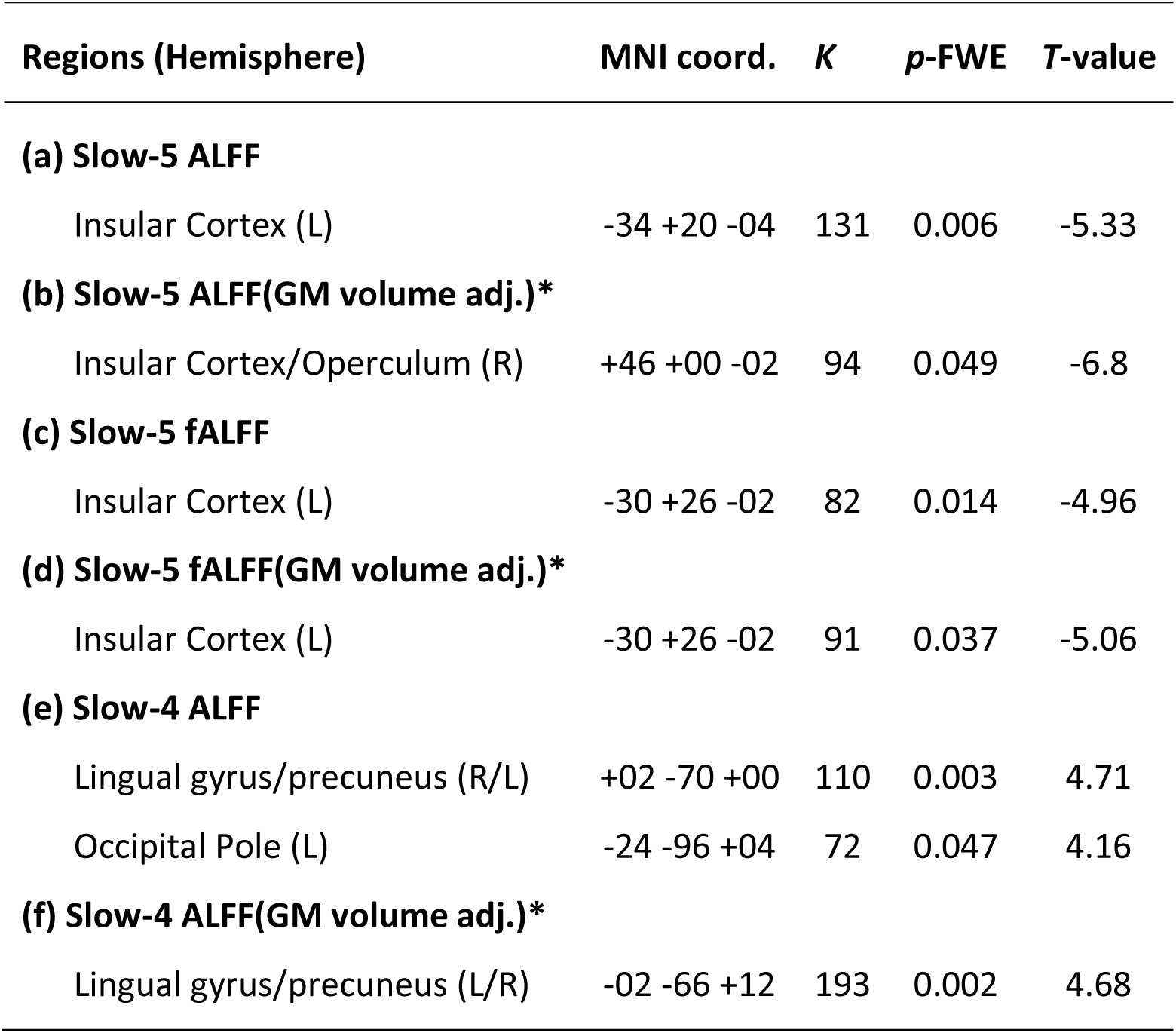
Regions demonstrating Drink x Age group interaction. Table displays regions that showed significant Drink x Age-group interaction (2-sided contrast): Peak regions, hemisphere, peak MNI coordinates (x, y, z), K= cluster size, p-values reported at p < 0.005 (height threshold) and cluster-level FWE corrected p < 0.05, T-value= peak of T-values. * = one-sided contrast.

| Regions (Hemisphere) | MNI coord. | K | p-FWE | T-value |
| --- | --- | --- | --- | --- |
| **(a) Slow-5 ALFF** |  |  |  |  |
| Insular Cortex (L) | -34 +20 -04 | 131 | 0.006 | -5.33 |
| **(b) Slow-5 ALFF(GM volume adj.)\*** |  |  |  |  |
| Insular Cortex/Operculum (R) | +46 +00 -02 | 94 | 0.049 | -6.8 |
| **(c) Slow-5 fALFF** |  |  |  |  |
| Insular Cortex (L) | -30 +26 -02 | 82 | 0.014 | -4.96 |
| **(d) Slow-5 fALFF(GM volume adj.)\*** |  |  |  |  |
| Insular Cortex (L) | -30 +26 -02 | 91 | 0.037 | -5.06 |
| **(e) Slow-4 ALFF** |  |  |  |  |
| Lingual gyrus/precuneus (R/L) | +02 -70 +00 | 110 | 0.003 | 4.71 |
| Occipital Pole (L) | -24 -96 +04 | 72 | 0.047 | 4.16 |
| **(f) Slow-4 ALFF(GM volume adj.)\*** |  |  |  |  |
| Lingual gyrus/precuneus (L/R) | -02 -66 +12 | 193 | 0.002 | 4.68 |

**Figure 3:**
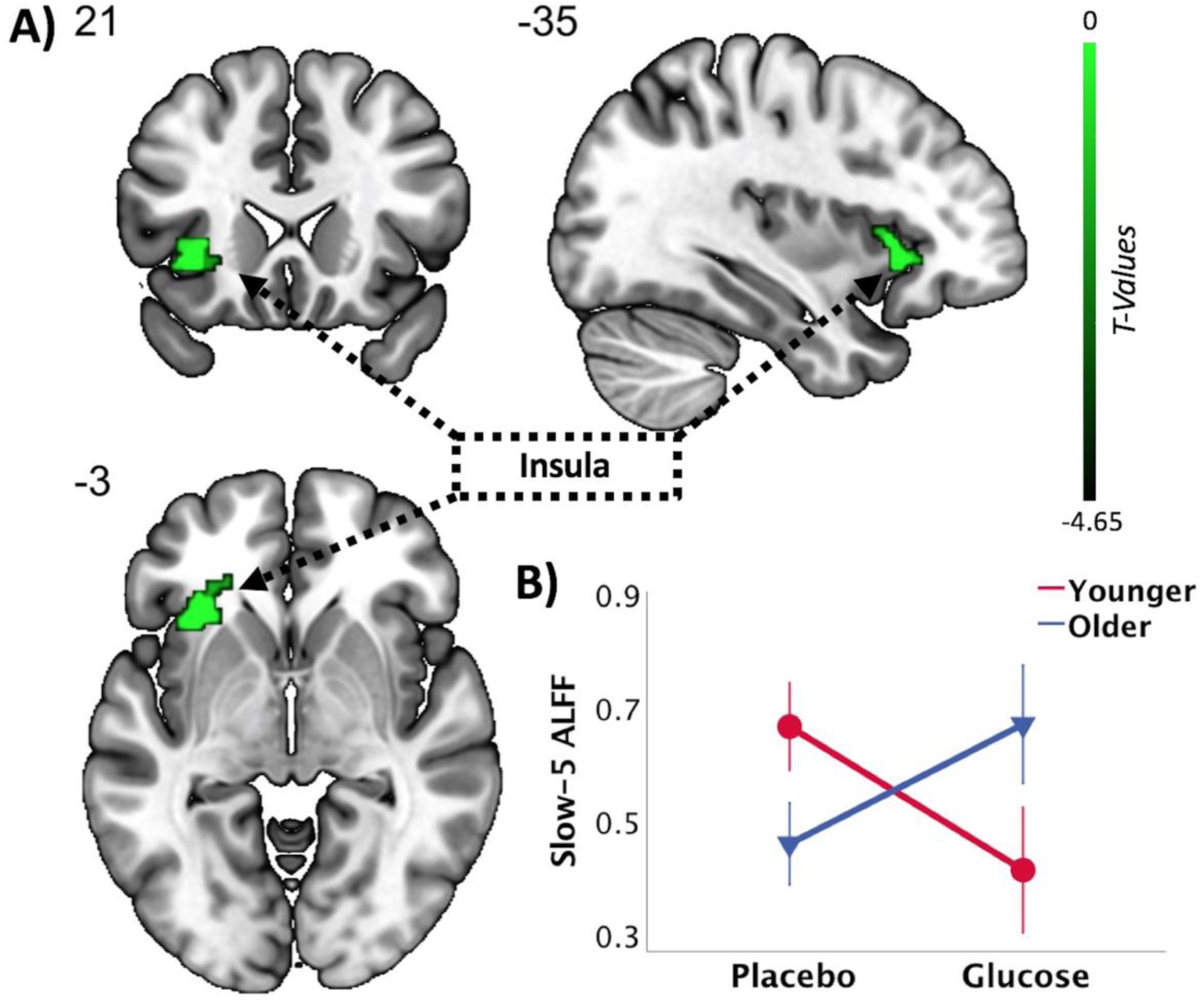
Slow-5 ALFF. (a) Drink x Age-group interaction in slow-5 (0.01-0.027 Hz) ALFF frequency band (young > older adults; Glucose > Placebo). Statistical images were assessed using a cluster-extend FWE corrected p-value < 0.05, obtained at a cluster-defining threshold of p< 0.005 (two-sided contrast). (b) Extracted slow-5 ALFF magnitude of left insular cortex cluster for each group per session (error bars reflect SEM).

Adjusting for GM volume did reveal a significant treatment x age group interaction in the opposite hemisphere in slow-5 ALFF (**Table 2b**), covering insular cortex and central operculum (Supplementary material S.2.1, Fig. 4).

**Figure 4:**
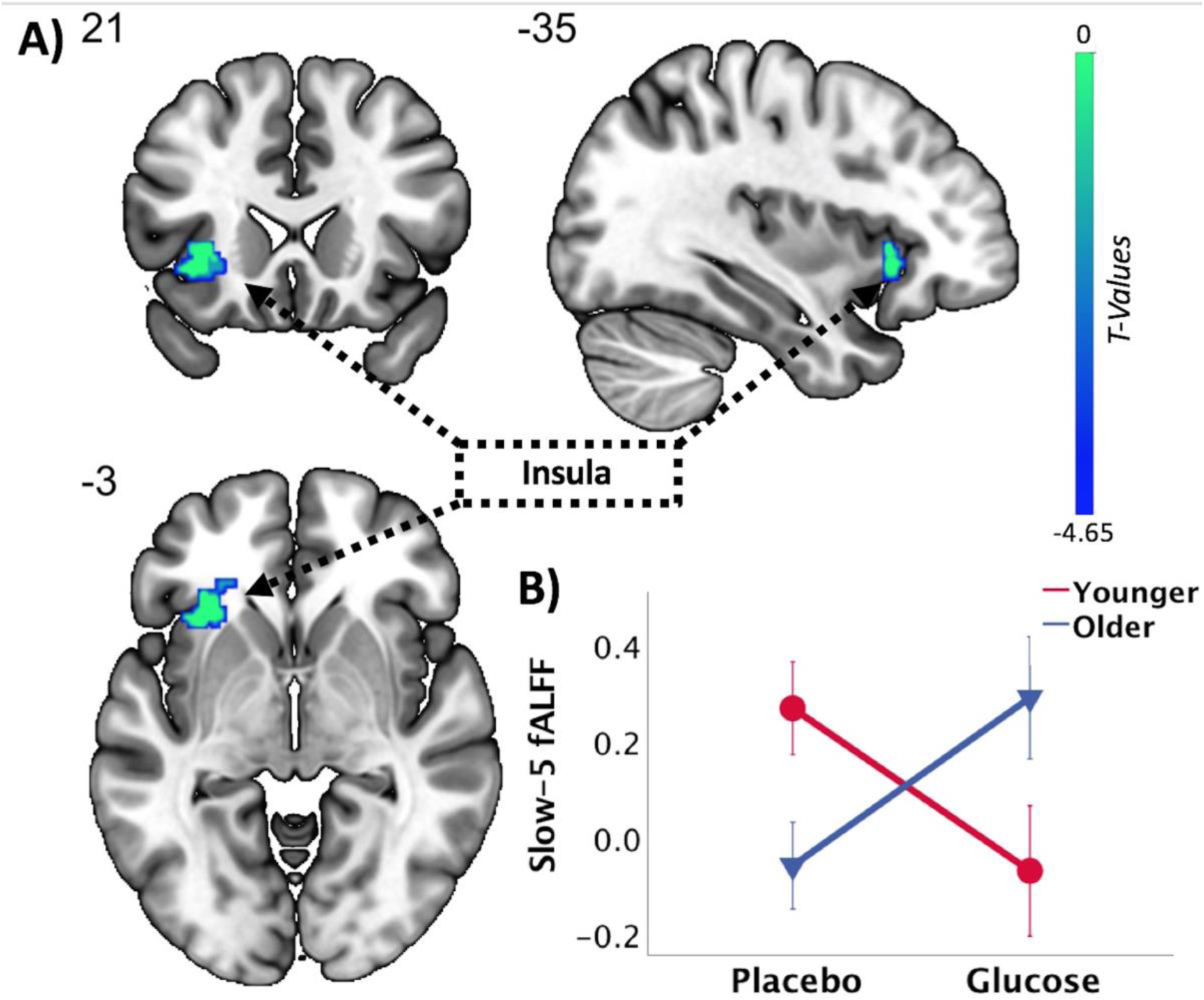
Slow-5 fALFF. (a) Drink x Age-group interaction in slow-5 (0.01-0.027 Hz) fALFF frequency band (young > older adults; Glucose > Placebo). Statistical images were assessed using a cluster-extend FWE corrected p-value < 0.05, obtained at a cluster-defining threshold of p < 0.005 (two-sided contrast). (b) Extracted slow-5 ALFF magnitude of left insular cortex cluster for each group per session (error bars reflect SEM).

#### Slow-5 (0.01-0.027Hz) fALFF

A significant Drink x Age-group interaction was evident in a cluster along the left insular cortex and frontal orbital cortex (FOrb) in the slow-5 fALFF (**Table 2c**; **Figure 4a**).

Similar to the results in the slow-5 ALFF analysis, slow-5 fALFF increased in the older participants after glucose ingestion, while slow-5 fALFF decreased in the younger group after glucose administration compared to placebo (**Figure 4b**). This cluster remained significant when controlling for grey matter volume (**Table 2d**).

#### Slow-4 (0.027-0.073Hz) ALFF: Treatment x age group interaction

In contrast to the pattern observed in the slow-5 frequency band, we observed a Drink x Age group interaction in a cluster ranging from lingual gyrus (LG) across intracalcarine cortex (ICC) to the precuneus, as well as in a cluster around the left occipital pole (OP)(**Table 2e**; **Figure 5a**). Slow-4 ALFF in the younger group increased after glucose ingestion in the cluster in the OP and cluster around lingual gyrus and precuneus, while a reduction in slow-4 ALFF in these clusters was observed in the older adults (**Figure 5 b&c**). Adjusting for grey matter volume resulted in a significant interaction in the cluster covering lingual gyrus and precuneus (**Table 2e**).

**Figure 5:**
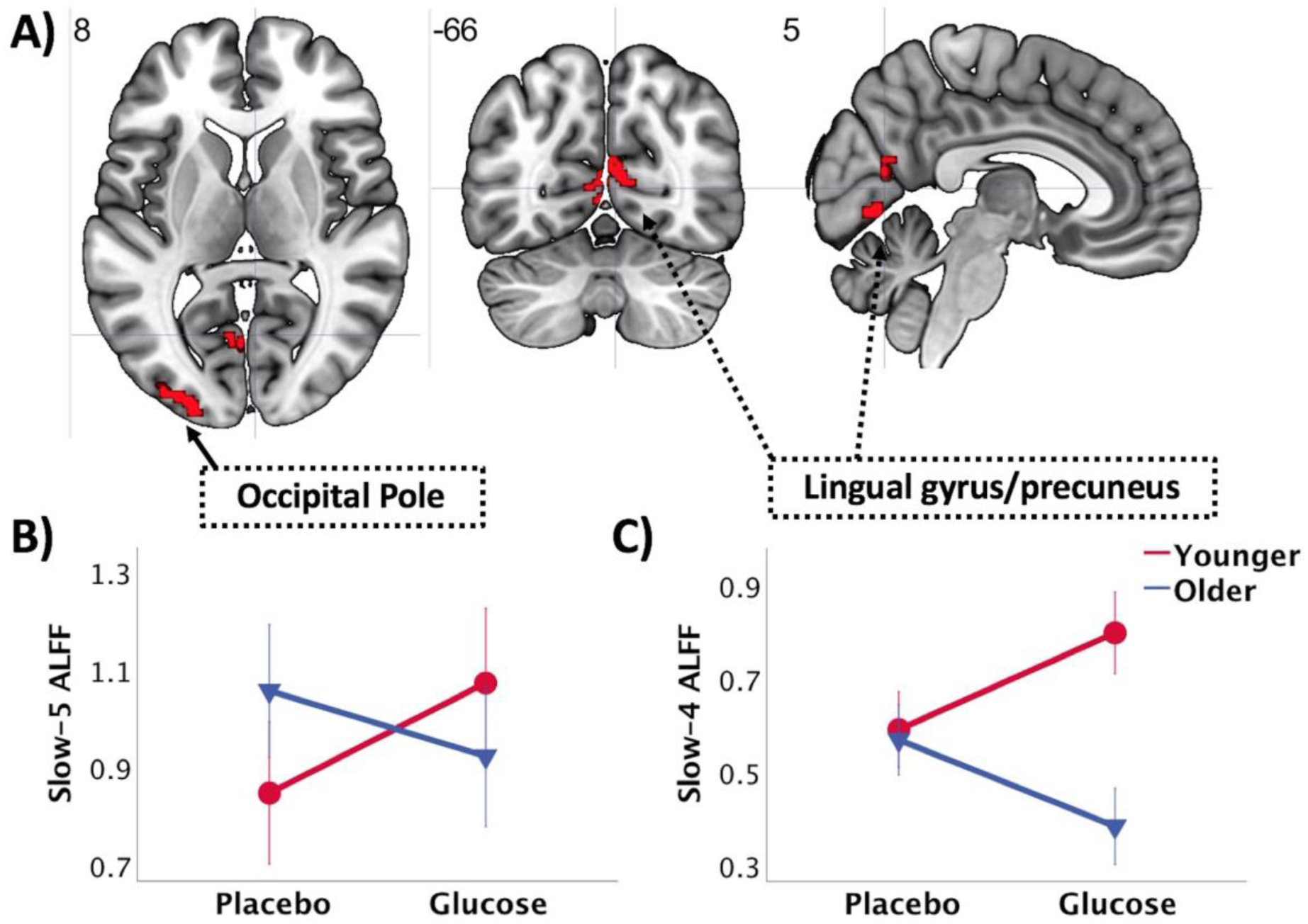
Slow-4 ALFF. (a) Drink x Age-group interaction in slow-4 (0.027-0.073Hz) ALFF frequency band (young > older adults; Glucose > Placebo). Statistical images were assessed using a cluster-extent FWE corrected p-value < 0.05, obtained at a cluster-defining threshold of p< 0.005 (two-sided contrast). (b) Extracted slow-4 ALFF magnitude of cluster in occipital pole for each group per session (error bars reflect SEM). (c) Extracted slow-4 ALFF magnitude of cluster in lingual gyrus/precuneus for each group per session (error bars reflect SEM)

#### Slow-4 (0.027-0.073Hz) fALFF

No significant Drink x Age-group interaction was observed in slow-4 fALFF.

## Discussion

We compared measures of brain activity in a group of younger and group of older adults on two occasions, once after the ingestion of a glucose drink and once after a placebo. We focused on a data-driven approach using frequency specific ALFF and fALFF subdivided into slow-5 (0.01-0.027Hz) and slow-4 (0.027-0.073Hz) frequency bands. We wanted to investigate frequency dependent changes in ALFF and fALFF in slow-5 and slow-4 frequency bands and to examine age-related abnormalities in the response to energy intake in these frequency bands.

### Slow-5 ALFF and fALFF

A cluster in the left insular cortex and frontal orbital cortex (FOrb) was affected differently in the slow-5 frequency window after glucose ingestion in younger compared to older adults measured by both ALFF and fALFF indices (**Figure 3 & 4).** Whereas the amplitude of the BOLD signal in the slow-5 frequency range decreased in young participants after glucose ingestion, it increased in older participants.

The insular cortex is one of the main areas in the brain associated with feeding behaviour (Wright et al., 2016). The anterior insular contains the primary gustatory cortex, containing neurons responding to different tastes and textures of food (Verhagen et al., 2004; Rolls, 2006). The insula together with the anterior cingulate cortex (ACC) comprise the primary components of the salience network (Menon and Uddin, 2010). In particular, the anterior part of insula has been suggested to be involved in a wide range of conditions and behaviours including the interoceptive representation of the physiological condition of the body (Craig and Craig, 2009). The insula has been implicated in the state of hunger, where hunger has been associated with increases in regional cerebral blood flow (rCBF), satiation has been associated with decreases in rCBF in the insula (Gautier et al., 2000).

The decrease in insular slow-5 ALFF/fALFF in the young participant group is consistent with other research in healthy young adults which has shown decreases in BOLD signal after the ingestion of glucose. Page et al (Page et al., 2013) reported reduced CBF in the insula amongst other regions (hypothalamus, thalamus, anterior cingulate and striatum) following glucose consumption. Further, a recent study reported decreased degree of centrality (DC) (a measure global connectivity of brain regions) in orbital inferior frontal gyrus and left anterior insula (Al-Zubaidi et al., 2018). An inverse relationship between levels of circulating blood glucose and left dorsal-insula response to food pictures has been reported (Simmons et al., 2013). Our results further support and extend these findings demonstrating that in healthy young adults, increasing circulating levels of blood glucose through oral administration resulted in a decrease in slow-5 ALFF and fALFF in the left insular cortex.

In the present study, older participants showed the opposite pattern of change in response to glucose ingestion than the younger group, an increase in ALFF and fALFF in the left insular cortex and orbitofrontal gyrus after glucose ingestion. The insula has been identified as a key region in the study of obesity and eating disorders. While a greater decrease in regional cerebral blood flow (rCBF) in the insular cortex has been reported in obese individuals after satiation (Gautier et al., 2000), abnormalities in insula activation have been reported in food cue paradigms in obese compared to lean subjects (Scharmüller et al., 2012), as well as aberrant functional connectivity in the resting state (Kullmann et al., 2012). Furthermore, the insula has been identified as a key region in the context of anorexia nervosa (AN) and bulimia nervosa, with dysfunction of the insula cortex regarded as a crucial risk factor for AN (Nunn et al., 2011).

Alterations in insula activity have also been reported in response to sweet tastes (Wagner et al., 2008) and studies using food cue paradigms (Ellison et al., 1998). Given the central role of the insular in hunger and satiety, and the abnormalities in signalling observed in neuroimaging studies in obesity and anorexia, our findings raise the suggestion that processing signals of energy intake in the insula are altered in aging, specifically in the slow-5 frequency range.

### Lateralisation of insula

We observed the group differences in response to glucose ingestion in the left insula. A recent meta-analysis reported greater involvement of the left insula in the context of appetite and food stimuli (Kelly et al., 2012). However, it should be noted that other meta-analyses reported greater involvement of the right insula (Kurth et al., 2010; Tang et al., 2012). It is noteworthy that when we controlled the slow-5 ALFF for GM, an interaction was observed in the right hemisphere, suggesting that subthreshold the effect might be observable bilaterally (Supplementary material Fig. 4).

### Slow-4 ALFF and fALFF

Slow-4 ALFF analysis revealed differences in young and older participants in response to glucose ingestion in the occipital pole and lingual gyrus and precuneus. In young adults an increase in BOLD amplitude was noted in these clusters, whereas a decrease in amplitude was observed in the older group. No significant interaction was observed in slow-4 fALFF. The precuneus is a hub in the default mode network (Cavanna and Trimble, 2006; Raichle, 2015), and involved in self-centred mental imagery strategies (Cavanna and Trimble, 2006). The DMN is active when participants are not focused on the external environment turn their attention inward (Buckner et al., 2008). Deactivation of the precuneus has been observed in response to satiation (Gautier et al., 2000), as well as to conscious suppression to food craving (Yokum and Stice, 2013). The deactivation observed in older participants could therefore be interpreted as a response to satiation.

### Slow-4 versus slow-5 ALFF and fALFF

It has been reported that slow-5 fluctuations are more dominant within prefrontal, parietal and occipital regions especially within components of the default mode network (DMN), while slow-4 fluctuations are more prominent in the basal ganglia (Zuo et al., 2010; Wang et al., 2016). Overall, our results suggest that ALFF and fALFF in the slow-5 frequency window may be linked with signalling related to energy homeostasis and could provide insights on normal and abnormal signalling in response to energy intake.

### Reported feelings of hunger

Unlike other studies that have shown that older people consistently rate themselves as less hungry compared to younger adults (Parker et al., 2004; Rolls et al., 1995), we did not observe any statistically significant differences in this study. However, mean values of ratings of hunger were still lower than those of younger adults (see table 3). Larger sample sizes might be warranted to see a significant effect.

### Limitations

We investigated the brain response to energy intake in a sample of healthy young and a group of healthy older adults who did not differ significantly in BMI. The older participants in this sample are therefore unlikely to show reduced body weight due to anorexia of ageing, however due to the cross-sectional nature of the study, this cannot be ascertained. We characterised age-related differences in response to glucose ingestion in healthy older adults,future studies should include participants with age related weight loss, to determine changes in brain activity show similar patterns.

Furthermore, we only included one VAS inquiring about how hungry participants were at different timepoints. For a more elaborate assessment of feelings related to hunger questions about fullness, desire to eat and question about prospective consumption can be included. Sex differences in response to taste in hunger and satiety conditions have been reported (Haase et al., 2011). Because of the limited sample size in the present study, we could not examine sex effects, future studies with bigger sample sizes are needed to replicate findings and examine potential sex related differences.

It may be argued that non-specific effects of aging were driving the observed effects, however, the fact that the present results were confined to anatomically relevant regions and all analyses were repeated controlling for grey matter volume, suggests this is a specific mechanism that was picked up.

## Conclusion

This is the first study to examine the differences in spontaneous neural activity in specific frequency bands following glucose ingestion in younger and older adults. Our results indicate that ingestion of glucose exerts different effects in young and older adults with respect to both brain loci and frequency. Changes were more consistent in slow-5 frequency band as they were observed in ALFF and fALFF. We conclude that activity in the insula, a key region involved in homeostatic energy balance, is differentially activated in younger and older adults following ingestion of glucose. Further research is suggested to identify differences in homeostatic brain networks to understand physiological changes that the brain undergoes in aging and to potentially identify targets for interventions against anorexia of aging.

## Acknowledgements

The study was partially supported by a scanning grant from Swinburne Neuroimaging (SNI) Facility, supported by the National Imaging Facility (NIF) under the National Collaborative Researcher Infrastructure Strategy (NCRIS).

