## Supplemental Material for "Resting state fMRI reveals differential effects of glucose administration on central appetite signalling in young and old adults"

### S.1 Results: Additional Visual Analogue Scales (VAS)

Subjective feelings of hunger, thirst, overall mood and attention were assessed on a computerized Visual Analogue Scale (VAS) before each blood sample. Participants were shown a straight 100 mm line with the endpoints labelled "not at all" on the left end and "extremely" on the right end, except for overall mood which was "extremely poor" on the left and "extremely good" on the right. They were instructed verbally to rate on the line how they felt in the present moment and to avoid the extreme ends of the line, as this would indicate the least and most they have ever felt in their life respectively.

Changes in VAS ratings from baseline to 20 min post-ingestion were assessed using repeated measures analysis of variance (rmANOVA) with two within group factors for treatment visit ("Drink": glucose, placebo) and timepoint sample taken ("Timepoint": baseline, 20mins post-ingestion) and between groups factor age group (young, old).

**Table 1: Table of means VAS Hunger**

| | | Placebo | | | | | | Glucose | | | | | |
|---|---|---|---|---|---|---|---|---|---|---|---|---|---|
| | | Young | | Old | | | | Young | | Old | | | |
| Rating | | Mean | SD | Mean | SD | T | P | Mean | SD | Mean | SD | T | P |
| **Fasting** | **Hunger** | 60.36 | 17.96 | 57.27 | 30.03 | 0.33 | 0.74 | 64 | 20.32 | 57 | 21.24 | 0.92 | 0.37 |
| **20min** | **Hunger** | 53.79 | 21.62 | 55.07 | 25.83 | -0.14 | 0.89 | 57.14 | 16.22 | 47.93 | 21.47 | 1.30 | 0.21 |
| **delta** | **Hunger** | -6.57 | 9.12 | -2.2 | 17.66 | -0.83 | 0.42 | -6.86 | 10.93 | -7.73 | 16.28 | 0.17 | 0.87 |

### S.1.1 Thirst

There was a significant Drink x Age Group interaction for VAS rating of Thirst ($F(1, 26)= 6.48$, $P= 0.017$).

**Figure 1:**

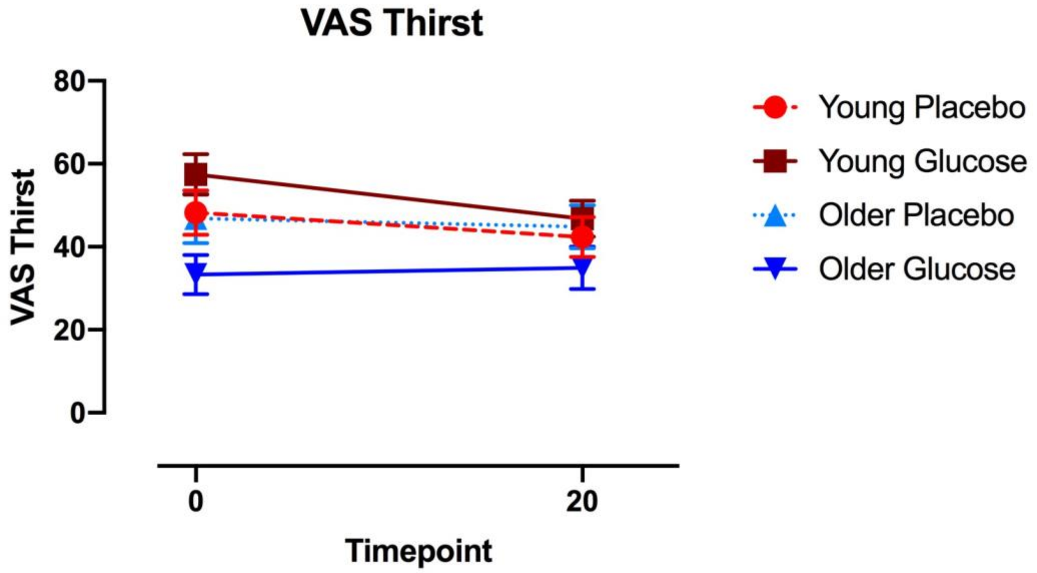


***Mean and SEM VAS Thirst ratings levels at baseline and 20 minutes post-dose dose at placebo and glucose visit for young (depicted in red) and older group (depicted in blue). Solid lines represent Glucose visit, dotted/dashed lines represent Placebo visit.***

**Table 2: Table of means VAS Thirst**

| | | Placebo | | | | | | Glucose | | | | | |
|---|---|---|---|---|---|---|---|---|---|---|---|---|---|
| | | Young | | Old | | | | Young | | Old | | | |
| **Rating** | | **Mean** | **SD** | **Mean** | **SD** | **T** | **P** | **Mean** | **SD** | **Mean** | **SD** | **T** | **P** |
| **Fasting** | **Thirst** | 48.21 | 20.17 | 46.87 | 23.05 | 0.167 | 0.87 | 57.43 | 18.37 | 33.31 | 18.74 | 3.549 | 0.001 |
| **20min** | **Thirst** | 42.36 | 17.87 | 44.8 | 20.26 | -0.343 | 0.73 | 46.79 | 16.28 | 34.93 | 19.64 | 1.762 | 0.089 |
| **delta** | **Thirst** | -5.86 | 10.52 | -2.07 | 14.84 | -0.79 | 0.44 | -10.64 | 16.35 | 2.73 | 19.75 | -1.98 | 0.06 |

### S.1.2 Overall Mood

There was a significant Timepoint x Age group interaction for VAS rating of overall mood (F(1,26)= 5.201, P= 0.031), a significant main effect of Timepoint (F(1, 26)= 11.89, P= 0.002) and a significant between subject effect (F(1, 26) -11.194, P= 0.003).

**Figure 2:**

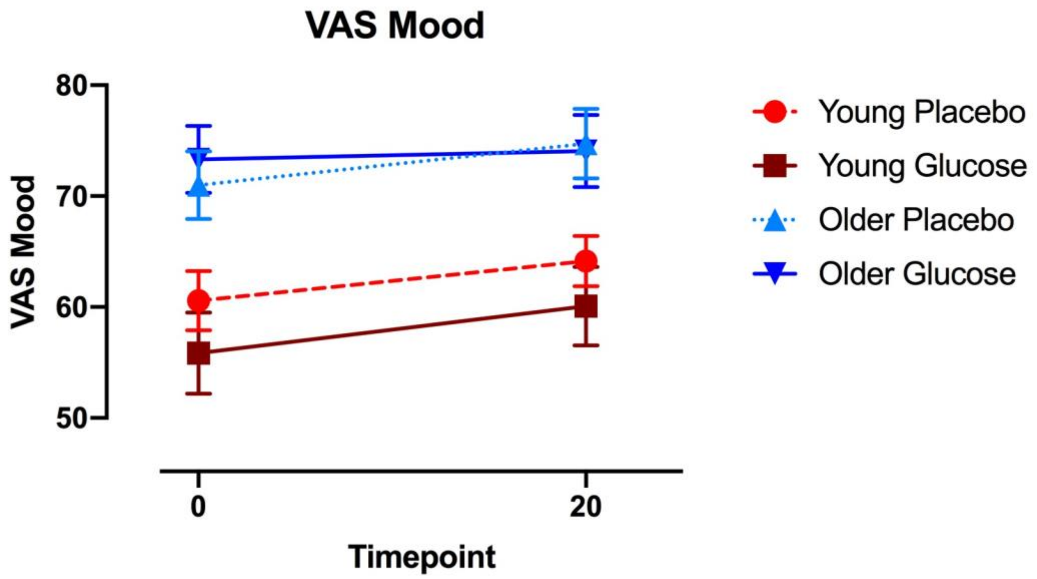


***Mean and SEM VAS Thirst ratings levels at baseline and 20 minutes post-dose dose at placebo and glucose visit for young (depicted in red) and older group (depicted in blue). Solid lines represent Glucose visit, dotted/dashed lines represent Placebo visit.***

**Table 3: Table of means VAS Mood**

| | | Placebo | | | | | | Glucose | | | | | |
|---|---|---|---|---|---|---|---|---|---|---|---|---|---|
| | | Young | | Old | | | | Young | | Old | | | |
| Rating | | Mean | SD | Mean | SD | T | P | Mean | SD | Mean | SD | T | P |
| Fasting | Mood | 60.57 | 10.03 | 71 | 11.874 | 12.595 | 0.017 | 55.86 | 13.722 | 73.31 | 12.037 | -3.713 | 0.001 |
| 20min | Mood | 64.14 | 8.51 | 74.73 | 12.19 | -2.69 | 0.01 | 60.07 | 13.29 | 74.07 | 12.60 | -2.91 | 0.01 |
| delta | Mood | 3.57 | 9.10 | 3.73 | 10.31 | 6.08 | 0.97 | 4.21 | 5.91 | 0.47 | 6.08 | 1.68 | 0.10 |

**S.1.3 Attention**

No interaction was observed for VAS rating of attention. However, there was a significant main effect of Timepoint (F(1,26)= 7.668, P= .01), as well as a between subject effect (F (1,26)= 12.327, P= 0.002).

**Figure 3:**

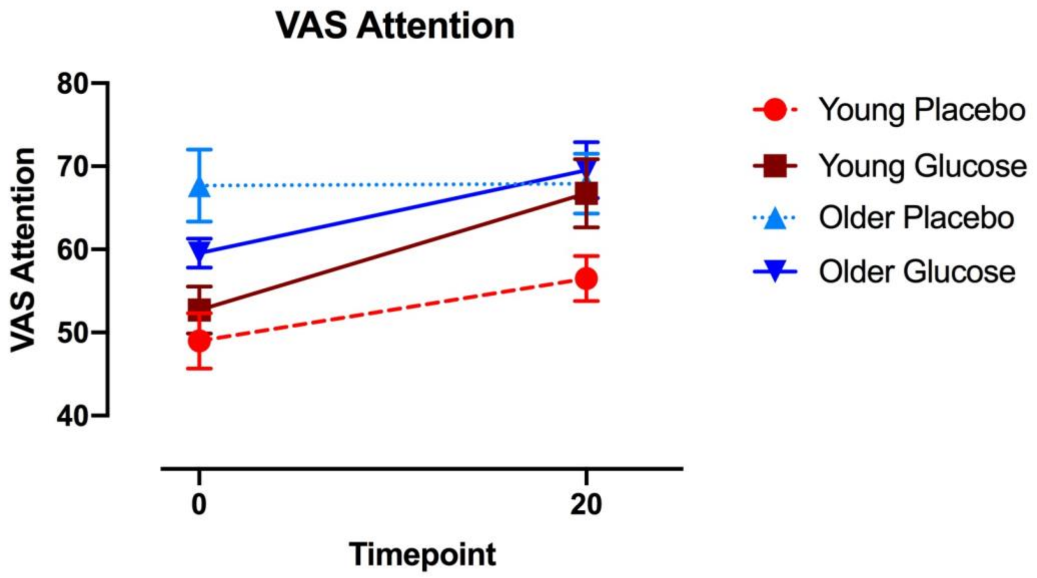


***Mean and SEM VAS Attention ratings levels at baseline and 20 minutes post-dose dose at placebo and glucose visit for young (depicted in red) and older group (depicted in blue). Solid lines represent Glucose visit, dotted/dashed lines represent Placebo visit.***

**Table 4: Table of means VAS Attention**

| | | Placebo | | | | | | Glucose | | | | | |
|---|---|---|---|---|---|---|---|---|---|---|---|---|---|
| | | Young | | Old | | | | Young | | Old | | | |
| Rating | | Mean | SD | Mean | SD | T | P | Mean | SD | Mean | SD | T | P |
| Fasting | Attention | 49.00 | 12.52 | 67.67 | 16.83 | -3.37 | 0.00 | 52.71 | 10.52 | 59.57 | 6.77 | -2.87 | 0.01 |
| 20min | Attention | 56.5 | 10.09 | 67.93 | 14.03 | -2.50 | 0.019 | 66.75 | 15.41 | 69.53 | 13.49 | -2.49 | 0.019 |
| delta | Attention | 7.5 | 8.91 | 0.27 | 13.33 | 1.705 | 0.1 | 6.86 | 8.80 | 0.47 | 8.40 | 2.00 | 0.055 |

### S.2 Overlap of frequency specific ALFF and fALFF results

#### S.2.1 Slow-5 Frequency band

**Figure 4:** **Slow- 5 ALFF and Slow-5 fALFF overlap**

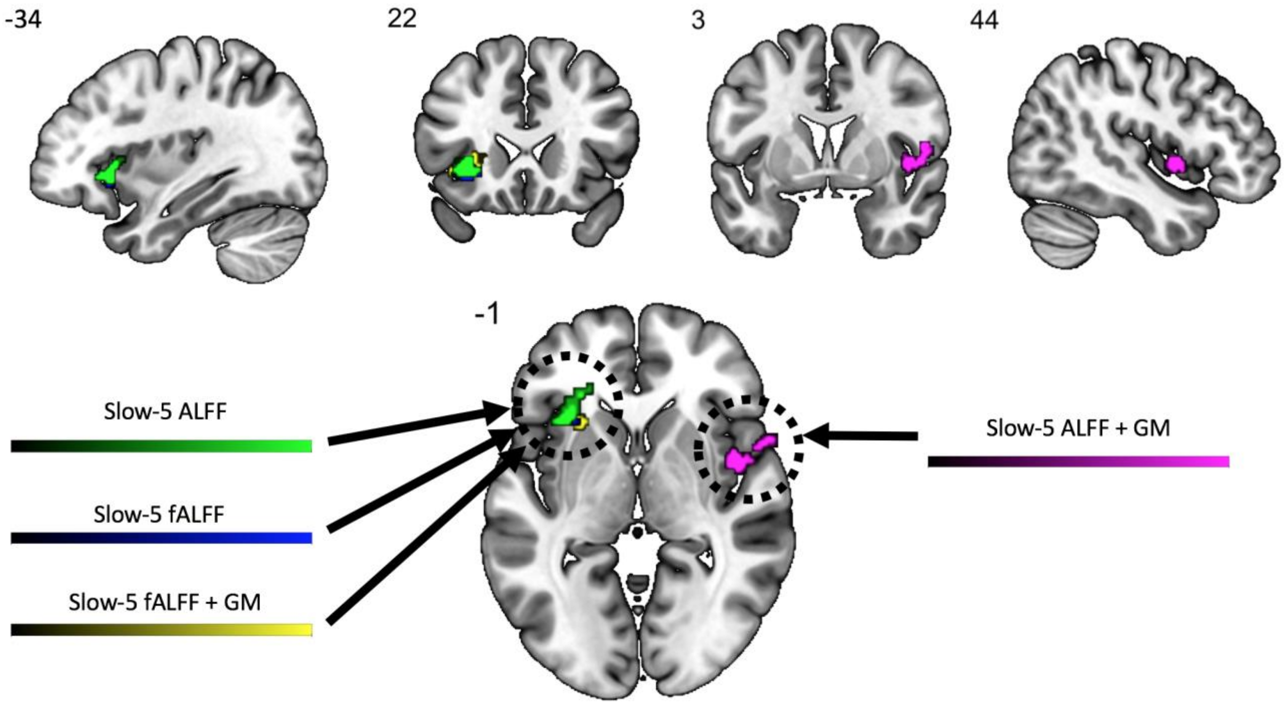


***Significant Drink x Age-group interactions from Slow-5 ALFF and fALFF analyses with and without covarying for gray matter.***

***Slow-5 ALFF results are depicted in green, slow-5 fALFF depicted in blue, slow-5 ALFF with covariate GM in purple, slow-5 fALFF with covariate GM in yellow.***

**S2.2 Slow-4 Frequency band**

**<u>Figure 5:</u> Slow-5 ALFF overlap**

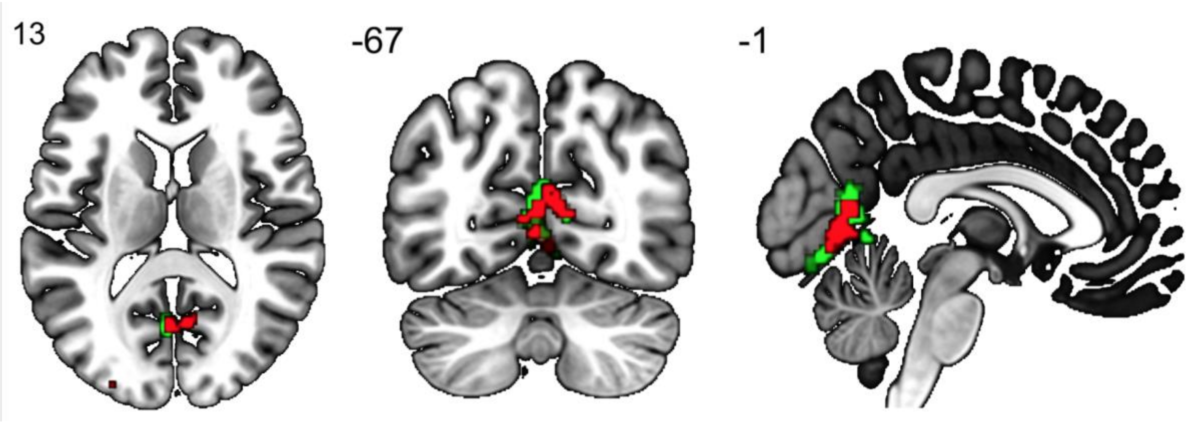


***Significant Drink x Age- group interaction in slow-4 ALFF with (depicted in green) and without (depicted in red) GM as covariate.***
